# Nuclear proteins acquiring Germinal Center B cells are specifically suppressed by follicular regulatory T cells

**DOI:** 10.1101/2022.10.25.513707

**Authors:** Fang Ke, Zachary L. Benet, Mitra P. Maz, Jianhua Liu, Alexander L. Dent, J. Michelle Kahlenberg, Irina L. Grigorova

## Abstract

Follicular regulatory T cells (Tfrs) restrict development of autoantibodies and autoimmunity while supporting high-affinity foreign antigen-specific humoral response. However, whether Tfrs can directly repress germinal center (GC) B cells that acquire auto-antigens is unclear. Moreover, TCR specificity of Tfrs to self-antigens is not known. Our study suggests that nuclear proteins contain antigens specific to Tfrs. Targeting of these proteins to antigen-specific B cells triggers rapid accumulation of Tfrs with immunosuppressive characteristics. Tfrs then exert negative regulation of GC B cells with predominant inhibition of the nuclear protein-acquiring GC B cells, suggesting an important role of direct cognate Tfr-GC B cells interactions for the control of effector B cell response.

## Introduction

Follicular regulatory T cells (Tfrs) are a subset of regulatory T cells, predominantly CXCR5^high^PD1^high^FoxP3^+^, that are present in B cell follicles of secondary lymphoid organs (SLOs) (Chung et al., 2011; Lim et al., 2004; Linterman et al., 2011; Wollenberg et al., 2011). Tfrs play multiple roles in the regulation of B cell responses. On one side, Tfrs prevent development of auto-Abs and autoimmunity (Botta et al., 2017; Clement et al., 2019; Fu et al., 2018; Gonzalez-Figueroa et al., 2021; Wu et al., 2016) and in some studies appear to have a modest negative effect on the GC and Ab response (Chung *et al*., 2011; Clement *et al*., 2019; Fu *et al*., 2018; Sage et al., 2016; Wollenberg *et al*., 2011). In addition to that, Tfrs affect B cell recruitment and selection in germinal centers (GCs) (Cavazzoni et al., 2022; Chung *et al*., 2011; Clement *et al*., 2019; Laidlaw et al., 2017; Lim *et al*., 2004; Linterman *et al*., 2011; Lu et al., 2021; Sage *et al*., 2016; Wollenberg *et al*., 2011; Wu *et al*., 2016; Xie et al., 2020).

GC B cell responses critically depend on the provision of help by Tfh cells. Tfh cells differentiate from foreign Ag-specific Th cells following immunization or infection. Importantly, differentiation of both Tfh and Tfr cells depends on the molecular interactions with antigen-presenting cells, involving SAP and ICOS molecules and the master transcription factor BCL6 (Hu et al., 2013; Johnston et al., 2009; Linterman *et al*., 2011; Nurieva et al., 2009; Pedros et al., 2016; Sage et al., 2013; Xu et al., 2013). Moreover, accumulation of Tfrs in SLOs occurs at the same time or slightly after the Tfh cells. However, in contrast to Tfh cells, the Tfr TCR repertoire predominantly overlaps with natural FoxP3^+^ Tregs and is more diverse than the Tfh TCR repertoire (Maceiras et al., 2017). While some studies suggest that TCR signaling must be important for Tfr development (Maceiras *et al*., 2017; Shrestha et al., 2015), which self-antigens are cognate to Treg-derived Tfrs is largely unknown. Such analysis is further complicated by very limited knowledge of the natural immunodominant self-antigens cognate to thymus-derived Tregs (Leonard et al., 2017). The deficiency in the identification of Tfr-specific self-antigens makes it difficult to assess whether Tfr may form direct cognate interactions with MHCII/self-peptides on the GC B cells.

Some experimental *in vivo* evidence supports the role of direct Tfr interactions with GC B cells for the negative regulation of immune responses. Tfr-produced membrane-attached neuritin has been recently shown to be acquired by B cells leading to their reduces ability to form plasma cells (Gonzalez-Figueroa *et al*., 2021). In addition, our previous study suggested that GC B cell-intrinsic production of CCL3 promotes direct contacts with Tfrs and modest inhibition of the foreign Ag-specific B cells at the peak of GC response (Benet et al., 2018). However, the contribution of direct Tfr-GC B cell encounters to the suppression of self antigen-acquiring and potentially autoreactive GC B cells is not known. The alternative hypothesis is that Tfrs may restrict autoreactive B cell responses non-specifically by reducing B cell activation/selection threshold through general inhibitory actions on B and/or Tfh cells.

In this study we show that nuclear proteins, often targeted by auto-Abs in autoimmune diseases, contain ligands that could trigger significant accumulation of Tfrs. Targeting these nuclear proteins to antigen-specific B cells by booster immunization induces rapid accumulation of Tfrs with elevated expression of immunosuppressive genes. Activated Tfrs then promote modest inhibition of the total GC responses with predominant suppression of the nuclear protein-acquiring GC B cells, memory B cells and plasma cells. These findings suggest an important role of direct Tfrs interactions with GC B cells presenting cognate to them self-antigens in the immunoregulation.

## Results

### Acquisition of Nuclear Proteins by GC B cells promotes rapid accumulation of Tfrs

Tfr deficiency leads to accumulation of anti-nuclear and tissue-specific antibodies (Abs) (Gonzalez-Figueroa *et al*., 2021), leading to development of auto-Ab-mediated autoimmunity in older mice (Fu *et al*., 2018). Nuclear proteins (NucPrs) that are often targeted in autoimmune diseases by auto-Abs include nucleosomal histones, SS-A/Ro, RNP-Sm (small nuclear ribonucleoproteins), Scl70 and Jo-1 (aminoacyl-tRNA synthetase, both nuclear and cytoplasmic) (Mahler and Fritzler, 2010; Tan, 1983). In this study we hypothesized that these NucPrs may contain peptides cognate to Tfr cells. Because the rise in Tfrs is usually observed at the same time or slightly after the increase in Tfh / GC B cells (Aloulou et al., 2016; Turner et al., 2017a), we first examined whether acquisition of NucPrs by GC B cells promotes accumulation of Tfrs (**Fig. 1A**). Bovine nucleosomes, SSA-Ro, RNP-Sm, Scl70 and Jo1 (that have over 92% homology to murine NucPrs) were biotinylated and conjugated to streptavidin to generate streptavidin-nuclear protein complexes (SA-NucPr). Alternatively, streptavidin was conjugated to foreign antigens (Ags): ovalbumin-biotin or duck egg lysozyme-biotin to generate SA-OVA or SA-DEL Ags respectively. C57BL/6 (B6) mice were subcutaneously immunized with SA-DEL to recruit both SA and DEL-specific B cells into the GC response. After formation of GCs (8 days later) mice were boosted with SA-NucPrs, SA or SA-OVA to promote acquisition of these Ags by the SA-specific GC B cells. The Ag-draining inguinal lymph nodes (dLNs) were collected 3 days later for flow cytometry analysis. We found that Tfr cells in the dLNs of SA-NucPrs-boosted mice were significantly increased as compared to mice boosted with SA or SA-OVA Ags (**Fig. 1A-C**). In contrast, the numbers of Tfh cells in the SA-NucPrs-boosted mice slightly decreased (**Fig. 1D**). The observed accumulation of CXCR5^high^PD1^high^FoxP3^+^ Tfr cells was not accompanied by increase in the frequencies of CXCR5^int^PD1^int^FoxP3^+^ or CXCR5^low^PD1^low^FoxP3^+^ Tregs (**Fig. 1B, E-G**). This data suggests that selected experimental scheme with the NucPrs promote rapid induction of Tfr cells in mice.

**Figure 1.**
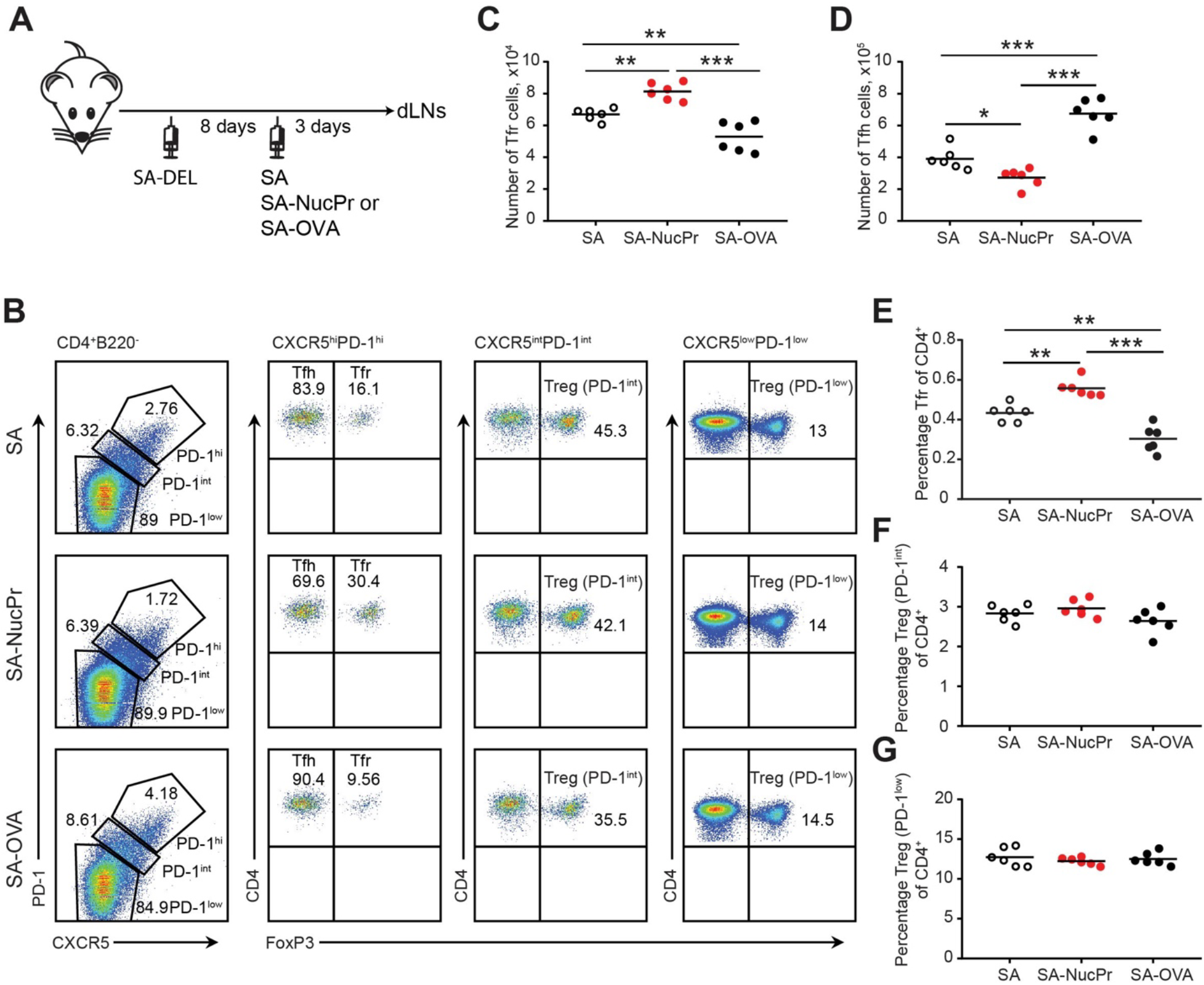
Boosting mice with SA linked to NucPrs induces rapid Tfr response. (**A**) Experimental outline. B6 mice were subcutaneously (s.c.) immunized with SA-DEL in Ribi and at day 8 were s.c. reimmunized with SA, SA-NucPr, or SA-OVA in Ribi for analysis 3 days later. (**B-G**) Flow cytometry analysis of Tfr, Tfh and other Treg subsets in the dLNs of immunized mice. (**B**) The gating strategy to identify the CXCR5^hi^PD1^hi^ FoxP3^+^ (Tfr) and CXCR5^hi^PD1^hi^ FoxP3^-^ (Tfh), CXCR5^int^PD1^int^ FoxP3^+^ (PD1^int^ Tregs) and CXCR5^low^PD1^low^ FoxP3^+^ (PD1^low^ Tregs) cell populations and representative flow plots for SA, SA-NucPr and SA-OVA boosted mice. (**C, D**) The numbers of Tfr cells (**C**) and Tfh cells (**D**). (**E-G**) The frequencies of Tfr cells (**E**), PD-1^int^ Tregs (**F**), and PD-1^low^ Tregs (**G**) of total CD4^+^ T cells in the dLNs. Data are representative of n=3 independent experiments. Each symbol represents one mouse. Lines indicate means. * *p*<0.05, ** *p*<0.01, *** *p*<0.001. One-way ANOVA analysis with Bonferroni’s multiple comparisons test.

### Nuclear Proteins promote accumulation of Tfrs with immunosuppressive gene expression profile

To assess gene expression and clonal repertoire of CXCR5^high^PD1^high^FoxP3^+^ Tfr cells induced by SA-NucPrs booster in mice, we performed 10x TCR immunorepertoire analysis of follicular (CXCR5^high^PD1^high^) T cells sorted from the dLNs of mice immunized with SA-DEL and boosted with SA or SA-NucPrs as described above (**Fig. 1A, Supplementary Fig. 1A**). Graph-based clustering analysis of follicular T cells revealed twelve clusters of cells that express Bcl6 with clusters 9-12 enriched for expression of FoxP3 (**Fig. 2A-D, Supplementary Fig. 1A-E, Supplementary Table 1**). Based on that and UMAP dimension reduction of the data, we will call clusters 9-12 as Tfr-like cells and clusters 1-8 as Tfh-like cells. Clusters 9 and 10 have majority of Tfr-like cells, while the cells in cluster 12 have upregulated expression of Mki67, Ccna2 and other genes associated with cell proliferation (**Fig. 2B-D, Supplementary Fig. 1D, E, Supplementary Table 1**). As previously reported (Maceiras *et al*., 2017), we found that TCR clonal repertoire of Tfr-like cells is very diverse, with majority of cells present at 1 cell per clone in the 10x data. We therefore performed analysis of the more abundant clones (>1 cell per clone) in **Fig. 2E, F**. Of note, the majority of TCR clones in clusters 9-12 overlapped with TCR clones from other follicular T cell clusters (**Fig. 2E**). In clusters 9 and 10 at least half of the clones were found exclusively in other Tfr-like clusters. In contrast, in clusters 11 and 12 the majority of clones were also present in Tfh-like clusters (**Fig. 2E, F**).

**Figure 2.**
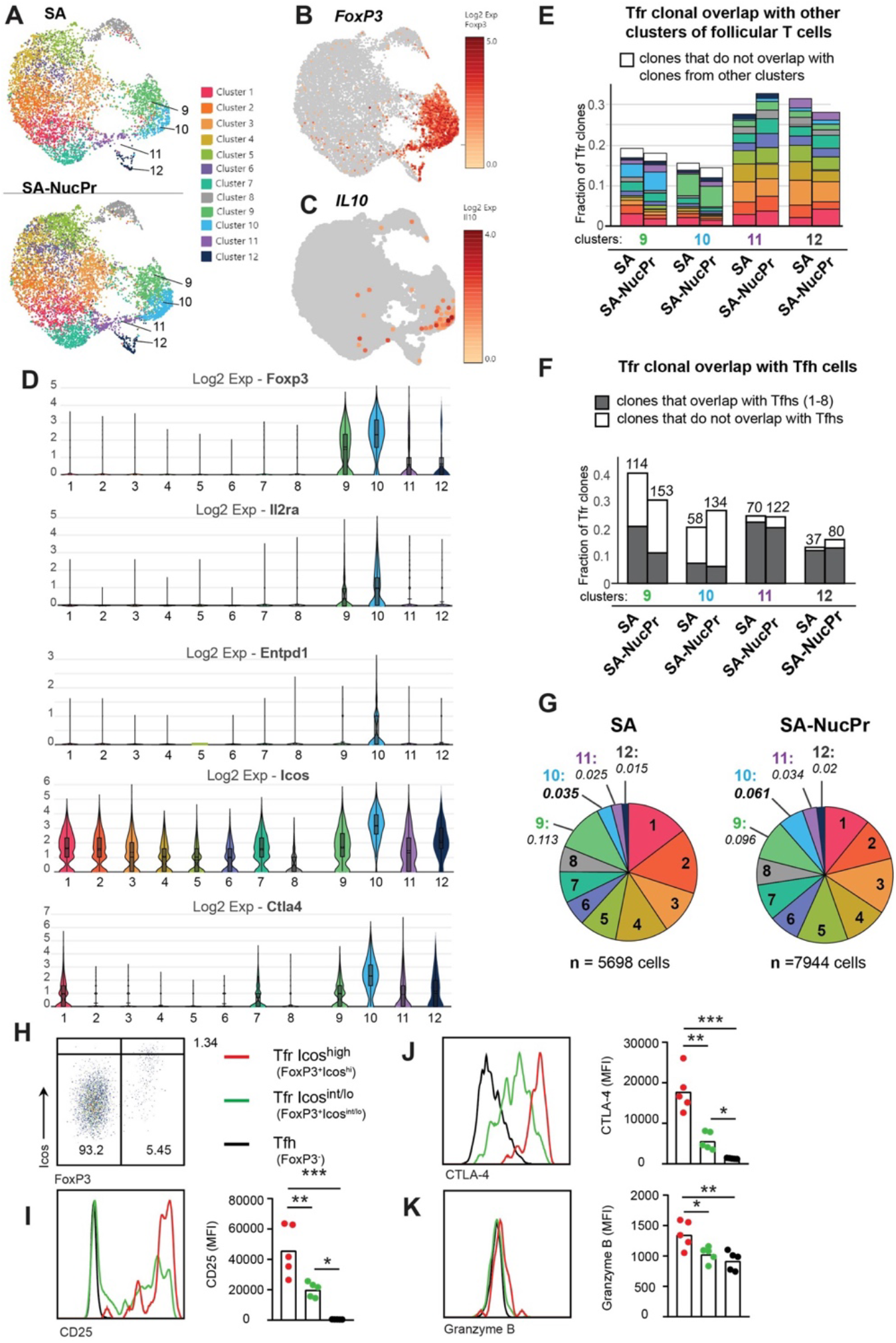
Analysis of Tfr cells gene expression and TCR repertoire in SA and SA-NucPr-boosted mice (related to Supplementary Fig. 1). **(A-G)** Single cell 10x genomics (Cell Ranger) analysis of gene expression and TCR repertoire of the CD4^+^CD8^-^220^-^CXCR5^hi^PD1^hi^ cells (Tfr and Tfh) sorted from the dLNs of mice immunized with SA-DEL and boosted with SA (2 mice) or SA-NucPr (2 mice) as in **Supplementary Fig. 1A**. (**A**) Graph-based clustering of follicular T cells from SA and SA-NucPr boosted mice visualized in 2D using uniform manifold approximation and projection for dimension reduction (UMAP) algorithm. (**B, C**) Single cell expression of FoxP3 (in **B**) and IL10 (in **C**) in follicular T cells for combined SA and SA-NucPr data shown in UMAP. (**D**) Expression of selected genes associated with Treg-mediated regulation in the follicular T cell clusters 1-12. Clusters 9-12 are enriched for follicular T cells with FoxP3 expression and will be called Tfr-like cell clusters. Clusters 1-8 will be called Tfh-like cell clusters. (**E**) Overlap of TCR clones (with > 1 cell per clone) within Tfr-like clusters 9-12 with other follicular T cell clusters (color-coded as in **2A**). Note that the largest fraction of TCR clones in cluster 9 overlap with cluster 10 and vice versa. (**F**) Fraction of TCR clones (with > 1 cell per clone) within Tfr-like clusters 9-12 that overlap or not with Tfh-like clusters 1-8 after SA vs. SA-NucPr boosting. Note that there is no increase in the Tfh-like TCR clones within Tfr-like clones after SA-NucPr boosting. (**G**) Relative abundance of follicular T cells in 1-12 clusters in the SA vs. SA-NucPr boosted mice. Note about 2-fold increase in the Tfr-like cluster 10 in the SA-NucPr boosted mice. (**H-K**) Flow cytometry analysis of CD25, CTLA4 and granzyme B expression in CD4^+^CD8^-^B220^-^ CXCR5^high^PD1^high^ cells that are FoxP3^-^ (Tfh), FoxP3+ Icos^high^ (Tfr Icos^high^) amd FoxP3 Icos^int/low^ (Tfr Icos^int/lo^) in the dLNs of SA-NucPr boosted mice. (**H**) The gating strategy of CD4^+^CD8^-^B220^-^ CXCR5^high^PD1^high^ cells. (**I-K**) Representative flow histograms (left panels) and MFIs (right panels) for CD25 (in **I**), CTLA4 (in **J**), Granzyme B (in **K**). n=2 independent experiments. Each point represents one mouse. * *p*<0.05, ** *p*<0.01, *** *p*<0.001. One-way ANOVA analysis with Bonferroni’s multiple comparisons test.

In parallel to the observed accumulation of CXCR5^high^PD1^high^FoxP3^+^ cells in mice boosted with SA-NucPrs by flow cytometry (**Fig. 1C, E**), 10x analysis revealed increase of cells in cluster 10 (**Fig. 2A, G**) that had the highest expression of FoxP3, ICOS, IL2ra, Entpd1, and CTLA4 (**Fig. 2D**). Cluster 10 was also enriched for cells expressing IL10 and granzyme B, while nrn1 (encoding Neuritin 1) - expressing cells were present in both clusters 9 and 10 and to some extent in other Tfr-like and Tfh-like cell clusters (**Fig. 2C, Supplementary Fig. 1F)**. In accord with the 10x data for cluster 10, flow cytometry analysis confirmed that CXCR5^high^PD1^high^FoxP3^+^ cells with the highest levels of ICOS also had elevated levels of CD25, CTLA4, and Granzyme B, as compared to Tfrs with ICOS^int/low^ and Tfh cells (**Fig. 2H-K)**. Overall, the data suggests that the accumulating Tfrs in SA-NucPr-boosted mice are likely to have more immunosuppressive Treg phenotype.

Importantly, we detected no increase in the Tfh-like cell TCR clones (from clusters 1-8) within Tfr-like clusters (9-12) in mice boosted with SA-NucPrs as compared to SA. Moreover, there was a trend for decreased frequency of Tfh-like cluster-associated clones among Tfrs in mice boosted with SA-NucPr (**Fig. 2E, F**). Therefore, based on the 10x TCR immunorepertoire analysis, we suggest that the observed increase in Tfrs in the SA-NucPrs boosted mice occurs due to accumulation of the cells with immunosuppressive phenotype that are not derived from abundant clones of Tfh cells.

### Tfr-like cells with immunosuppressive gene expression profile in human lymph nodes

To determine whether human Tfrs have any subsets similar to the murine Tfrs with immunosuppressive phenotype, we performed PCA analysis of the publicly available integrated multimodal single-cell data from human LN samples (GSE195673). We performed graph-based clustering analysis of CD4 T cells and identified follicular T cell clusters using a modular score generated based on the expression of CXCR5, PDCD1, BCL6, ICOS, CTLA4, IL1R2 and CXCL13 (**Supplementary Fig. 2A**). Follicular T cell subsets were then reclustered and expression of FoxP3 and other genes associated with Treg’s immunosuppressive activity were analyzed (**Supplementary Fig. 2A-C**). Within 8 of the detected follicular CD4 T cell clusters, FoxP3 expression was detected in clusters 4-7, with the highest expression in cluster 5 and the lowest in cluster 7. Elevated expression of IL2RA and ENTPD1, CTLA4, LAG3 was detected in clusters 5-7. Expression of TGFbetta1 and ICOS was the highest in cluster 7. Importantly, clusters 6 and 7 were also enriched in IL10-expressing cells (**Supplementary Fig. 2B, C**). Therefore, based on the single cell analysis, human LNs have Tfr-like cells that resemble gene expression characteristics of the immunosuppressive murine Tfr subset (cluster 10), but may have reduced or undetectable expression of FoxP3.

### Tfrs specifically suppress GC B cells acquiring nuclear proteins and exert more modest inhibition of total GC B cells

To determine whether NucPrs-induced Tfrs can in turn affect GC responses, we examined the total and SA-specific GC B cells in the SA-DEL-immunized mice boosted with SA-NucPrs or control Ags (**Fig. 1A)**. We found that in mice boosted with SA-NucPrs the frequency of GC B cells was reduced (**Fig. 3A, B, Supplementary Fig. 3A**). Moreover, SA-specific GC B cells that were expected to preferentially reacquire SA-linked NucPrs, underwent greater suppression than total GC B cells or DEL-specific B cells (**Fig. 3A, C, D**; **Supplementary Fig. 3B-F**). Of note, in mice boosted with SA-OVA no decrease in the total or SA-specific GC B cells was detected (**Fig. 3A-D**). In parallel to the observed reduction in the numbers of SA-specific GC B cells, we detected reduced accumulation of the SA-specific class-switched memory B cells in mice boosted with SA-NucPrs (**Fig. 3E-G)**. We also found decreased formation of SA-specific plasmablasts (PB), including the PB recently derived from GC B cells (GCPB) (**Fig. 3H-L**). Finally, in mice boosted with SA-NucPrs the titers of the SA-specific Abs were reduced as compared to mice boosted with SA (**Supplementary Fig. 3G)**.

**Figure 3.**
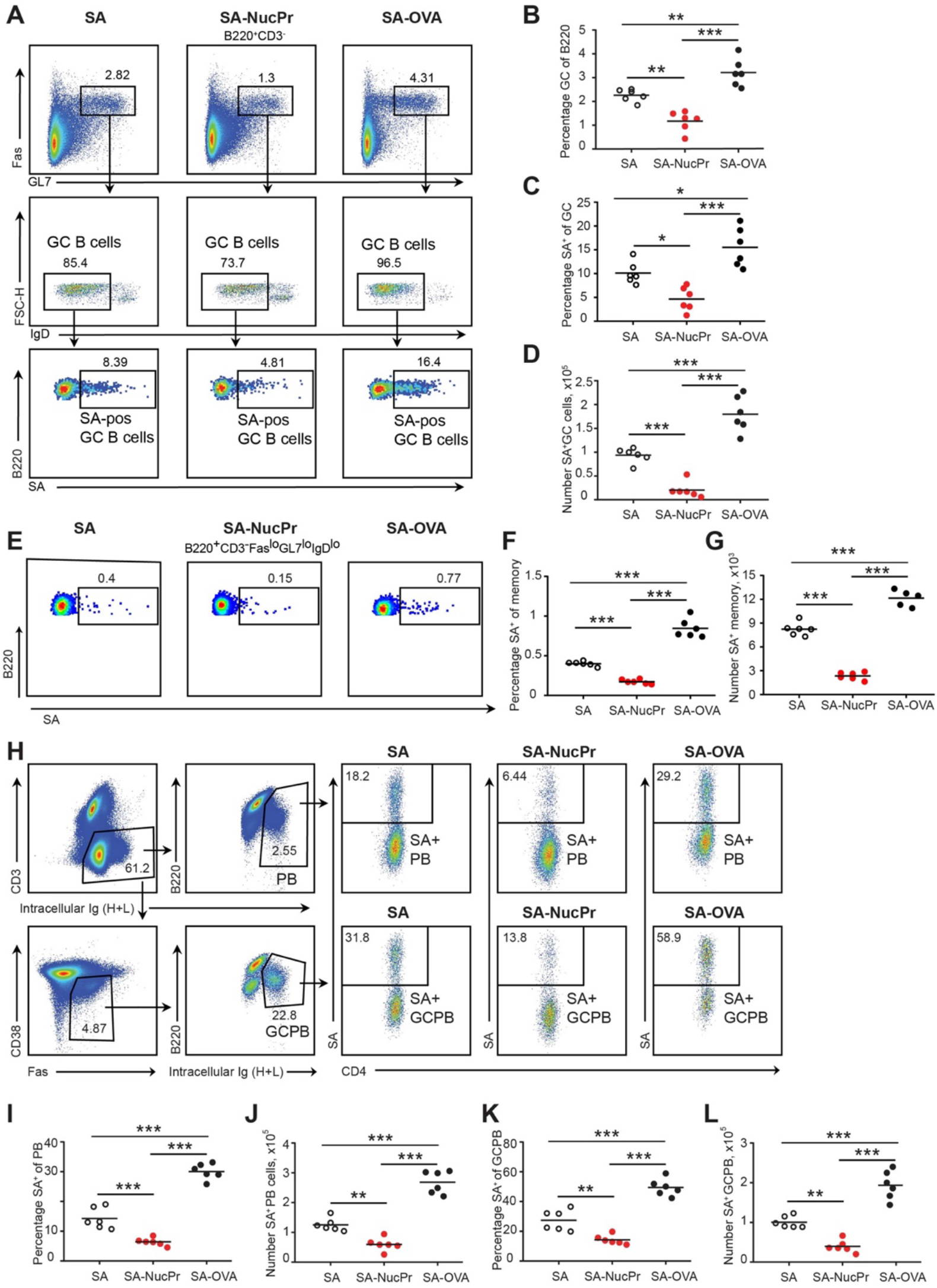
Boosting mice with SA-NucPrs suppresses GC, memory and PB responses with predominant inhibition of the SA-specific B cells (related to Supplementary Fig. 3). Flow cytometry analysis of the total and streptavidin (SA)-specific GC (**A-D**), memory (**E-G**) and PB (**H-L**) responses in the dLNs of mice treated as shown in **Fig. 1A**. (**A**) Representative flow plots for the total and SA-specific GC B cells in SA, SA-NucPr and SA-OVA boosted mice. (**B**) The GC B cells percentage of total B220^+^ B cells. (**C, D**) SA-specific GC B cells percentage of total GC B cells (**C**) and their numbers in dLNs (**D**). (**E**) Representative flow plots for the class-switched SA-specific memory B cell response in SA, SA-NucPr and SA-OVA boosted mice. (**F, G**) SA-specific memory B cells percentage of total class-switched memory B cells (B220^+^CD3^-^FAS^lo^GL7^lo^IgD^lo^) (**F**) and their numbers in dLNs (**G**). (**H)** Representative flow plots for SA-specific PB and GCPB response in SA, SA-NucPr and SA-OVA boosted mice. (**I-L**) SA-specific PB and GCPB percentage of total PB and GCPB (**I, K**) and their numbers in dLNs (**J, L**). Data are representative of n=3 independent experiments. Each symbol represents one mouse. Lines indicate means. * *p*<0.05, ** *p*<0.01, *** *p*<0.001. One-way ANOVA analysis with Bonferroni’s multiple comparisons test.

Importantly, our suggested immunization scheme (**Fig. 1A**) promoted suppression of the NucPrs- acquiring GC B cells not only in the B6 mice but also in NZM2328 mice. NZM2328 mice spontaneously develop dsDNA auto-Abs and systemic lupus erythematosus (SLE)-like disease after 5 months of age (Rudofsky and Lawrence, 1999; Waters et al., 2001; Wolf et al., 2018) and form anti-ribonucleoprotein (RNP) auto-Abs (**Fig. 4A)**. In this study we found that the ratio of GC B cells to Tfh cells (that is highly conserved in GCs (Baumjohann et al., 2013)) is elevated over 5 fold in 6-8 week old female NZM2328 compared to B6 mice, suggesting dysregulated GC responses even in young NZM2328 mice (**Fig. 4B, C**). Importantly, SA-DEL immunization followed by boosting with SA-NucPrs decreased the GC B/Tfh cell ratio in the NZM2328 mice, as well as in B6 mice (**Fig. 4B, C**). Even more significant was the observed decrease in the SA- specific GC B cell/Tfh cell ratio in the SA-NucPrs boosted NZM2328 (**Fig. 4D, E**). Therefore, even in pre-autoimmune state in mice where B cell responses are inflated, SA-DEL immunization followed by SA-NucPrs boosting provided a suppressive effect.

**Figure 4.**
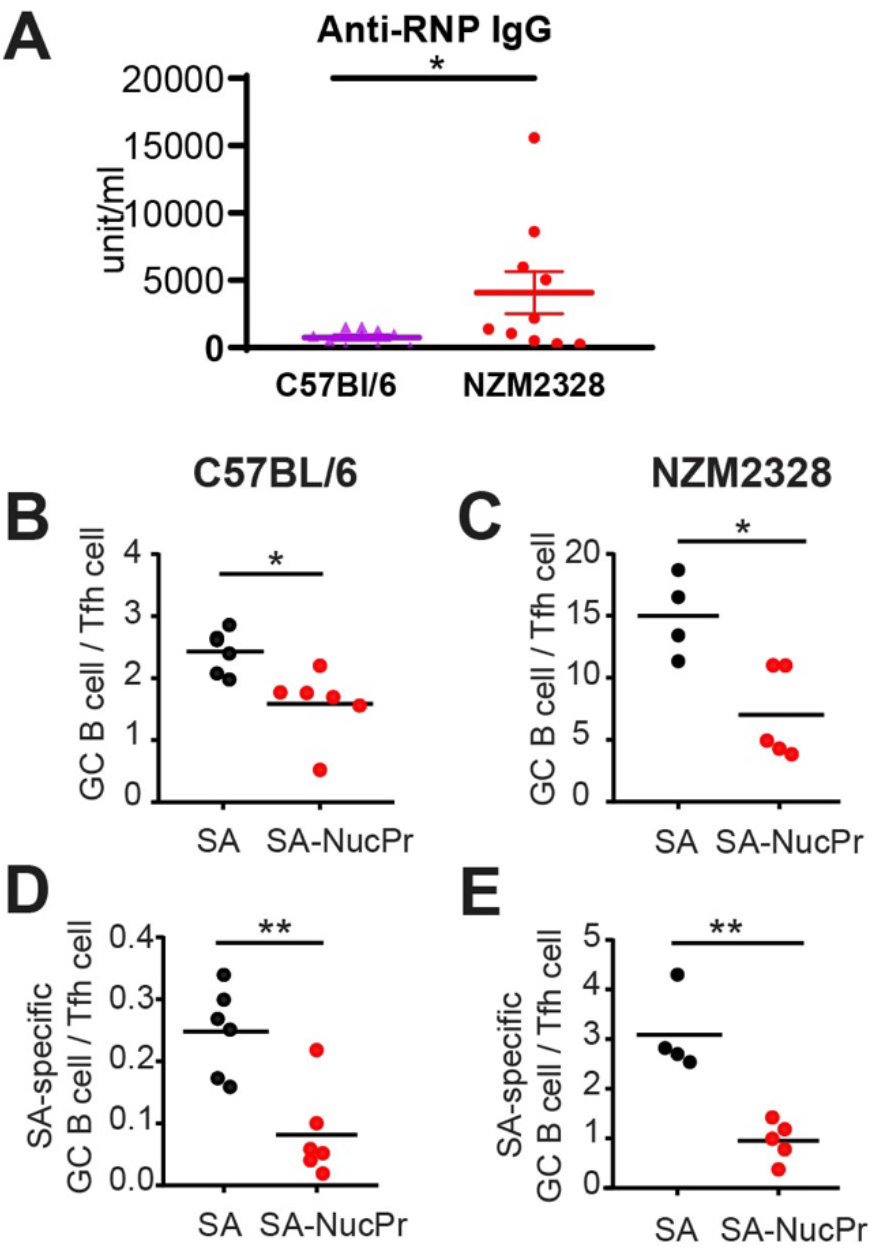
Boosting mice with SA-NucPrs reduces the ratio of GC B cells to Tfh cells in both C57BL/6 and NZM2328 mice. (**A**) Analysis of anti-RNP Abs via ELISA in the serum of female C57BL/6 (52 weeks old) and NZM2328 female mice (32-52 weeks old). (**B-E**) Suppression of GC B cell responses in the dLNs of 6-8 weeks old C57BL/6 and NZM2328 female mice treated as in **Fig. 1A** after the boost with SA-NucPr. The ratio of GC B cells to Tfh cells in C57BL/6 (**B**) and NZM2328 (**C**) mice. The ratio of SA-specific GC B cells to Tfh cell in in C57BL/6 (**D**) and NZM2328 (**E**) mice. n=2 independent experiments. Each symbol represents one mouse. Lines indicate means. * *p*<0.05, ** *p*<0.01. Two-tailed Student’s *t* test.

To verify whether observed suppression of the GC responses in the SA-NucPrs-boosted mice depended on Tfr cells, we repeated the immunization/ boosting experiments (as in **Fig. 1A**) in FoxP3-cre Bcl6^fl/fl^ mice that are deficient for Tfrs (Wu *et al*., 2016) or in control FoxP3-cre Bcl6^+/+^ mice. As expected, the frequency of CXCR5^high^ PD1^high^ FoxP3^+^ Tfrs in the FoxP3-cre Bcl6^fl/fl^ mice was low, and no increase in Tfrs was observed after boosting these mice with SA-NucPrs (as opposed to FoxP3-cre Bcl6^+/+^ control mice) (**Fig. 5A-C**). Importantly, while boosting FoxP3-cre Bcl6^+/+^mice with SA-NucPrs promoted significant suppression of the SA-specific and total GC B cells, both effects were completely abolished in the Tfr-deficient FoxP3-cre Bcl6^fl/fl^ mice (**Fig. 5D-G**). This data suggests a direct role of NucPrs-induced Tfrs in the negative regulation of the GC-responses with predominant suppression of the NucPrs-acquiring GC B cells.

**Figure 5.**
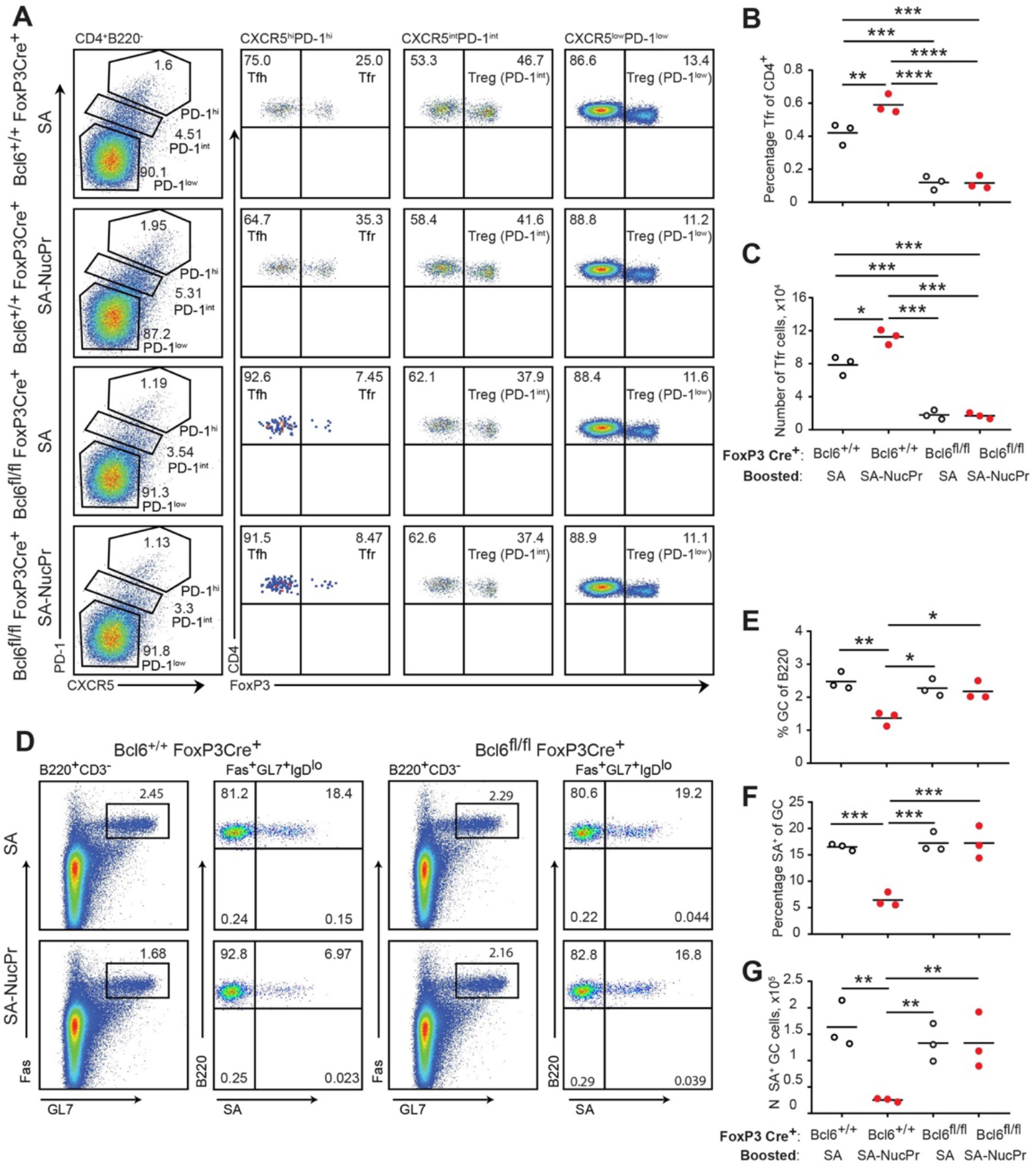
Tfrs are required for the suppression of GC response in mice boosted with SA-NucPrs. Bcl6^+/+^FoxP3Cre^+^ and Tfr-deficient Bcl6^fl/fl^FoxP3Cre^+^ mice were s.c. immunized with SA-DEL in Ribi and at day 8 reimmunized with SA, or SA-NucPr in Ribi s.c. for analysis 3 days later. (**A**) Representative flow plots for Tfr and other Treg subsets in the dLNs of mice boosted with SA or SA-NucPr. (**B, C**) Tfr cells percentage of CD4^+^ T cells (**B**) and total numbers (**C**) in dLNs. (**D**) Representative flow plots showing GC B cells and SA-specific GC B cells in Tfr-proficient and deficient mice after the boost with SA or SA-NucPr. (**E**) GC B cells percentage of total B220^+^ B cells. (**F, G**) SA-specific GC B cells percentage of total GC B cells (**F**) and total numbers (**G**) in dLNs. n=3 independent experiments. Each symbol represents one mouse. Lines indicate means. * *p*<0.05, ** *p*<0.01, *** *p*<0.001, **** *p*<0.0001, Two-way ANOVA analysis with Tukey’s multiple comparisons test.

### Identification of the immunization strategy that triggers rapid accumulation of Tfrs and suppression of GC response

We then examined which immunization strategy promotes accumulation of the NucPrs-induced Tfr cells (**Fig. 6**). Direct immunization of naïve mice with SA-NucPrs did not induce rapid expansion of Tfrs that was observed in mice preimmunized with SA-DEL (**Fig. 6A-D, left panels**). We then assessed whether anti-SA Abs (that should be induced after preimmunization of mice with SA-DEL) may facilitate SA-NucPrs uptake, presentation and efficient activation of Tfrs by antigen-presenting cells. To address this, naive mice were transferred with serum from SA-immunized mice and then immunized with SA-NucPrs. However, at 3 days following vaccination no effect on the Tfr cell frequency was detected as compared to mice that did not get serum transfer (**Fig. 6A, C, E, left panels**). We next examined whether accumulation of SA-specific B cells induced by preimmunization of mice with SA-DEL was important for rapid increase in Tfrs. To test this, we compared induction of the Tfr response by SA-NucPrs boosting in mice preimmunized with SA-DEL or with OVA. In contrast to the rapid expansion of Tfrs and suppression of GCs in the SA-DEL-preimmunized mice, no increase in Tfrs and/or suppression of GC B cells was detected in mice preimmunized with OVA (**Fig. 6B, D, F**). Since boosting of mice with an ongoing SA-specific GC response with SA-NucPrs is expected to promote acquisition of these Ags by the SA-specific GC B cells, we suggest that directing a combination of NucPrs to a selected subset of GC B cells may be a quick and efficient method for triggering rapid induction of Tfrs and suppression of GC B cell subset that present the acquired NucPrs’ peptides.

**Figure 6.**
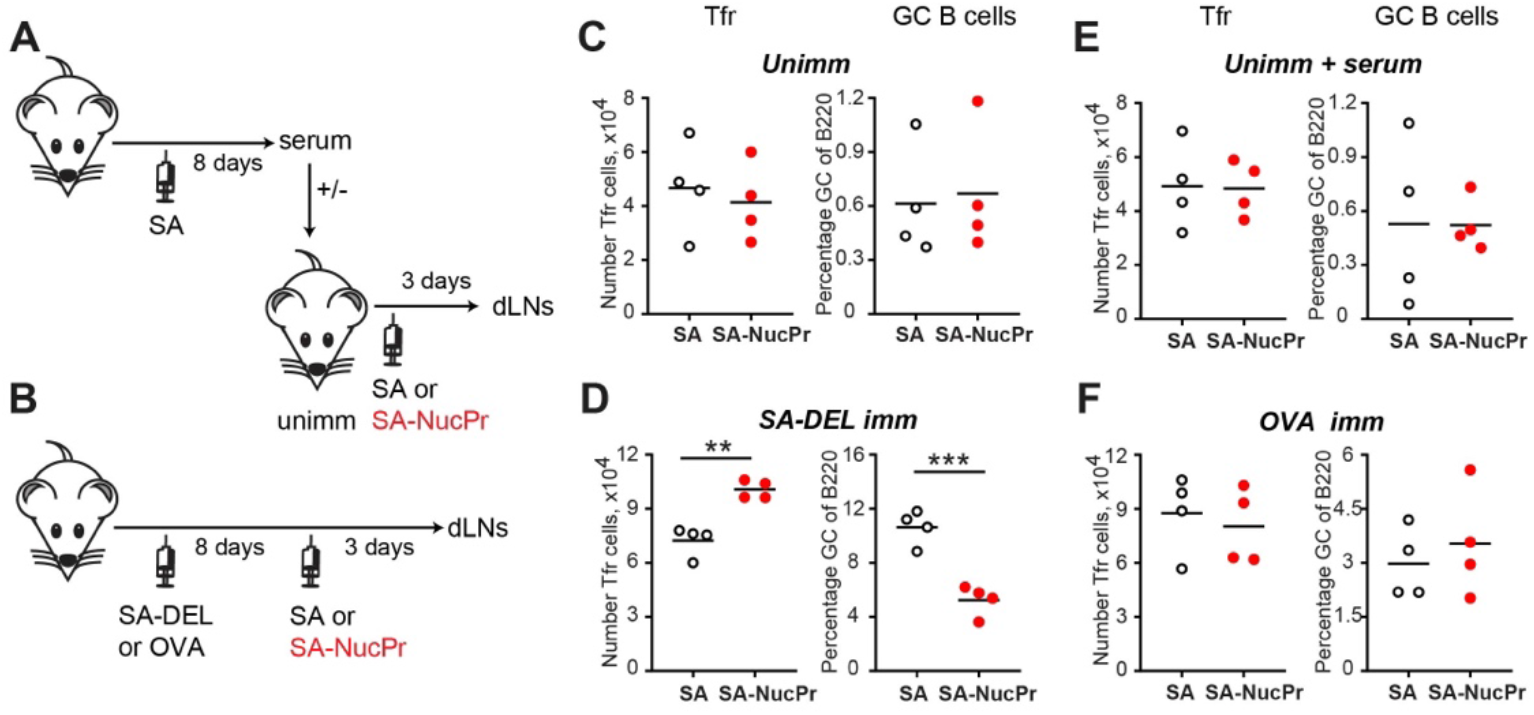
Analysis of the immunization conditions inducing rapid Tfr response. (**A, C, E**) Experimental outline (**A**). Some B6 mice were s.c. immunized with SA and serum were collected at day 8. Unimmunized B6 mice were transferred with serum from above (**E**) or not (**C**) and were s.c. immunized with SA or SA-NucPr in Ribi for analysis 3 days later. (**B, D, F**) Experimental outline (**B**). B6 mice were s.c. immunized with SA-DEL (**D**) or OVA (**F**) in Ribi and at day 8 were s.c. boosted with SA or SA-NucPr in Ribi for analysis 3 days later. (**C, D, E, F**) The numbers of Tfr cells (left panels) and the GC B cells percentages of total B220^+^ B cells (right panels). Data are from n=2 independent experiments. Each symbol represents one mouse. Lines indicate means. ** *p*<0.01, *** *p*<0.001, Two-tailed Student’s *t* test.

We then examined which of the selected NucPr Ags (nucleosomes, SSA-Ro, RNP-Sm, Scl70 and Jo1) promotes the observed accumulation of Tfrs and suppression of GC B cells. To address this, SA-DEL preimmunized mice were boosted with SA conjugated to each one of these Ags to assess Tfr cell frequency and SA-specific GC response (**Supplementary Fig. 4A-D**). While robust increase in Tfrs was detected in mice boosted with combined SA-NucPrs (**Supplementary Fig. 4B**), boosting with separate SA-NucPr Ags did not promote significant increase in Tfr cells (**Supplementary Fig. 4C**). Based on this data we suggest that the observed increase in Tfrs (**Supplementary Fig. 4B**) occurs due to additive accumulation of Tfrs specific to various NucPrs Ags.

Despite of the non-significant increase in Tfrs after targeting separate NucPrs to SA-specific B cells, we detected decreased frequency of the SA-specific GCs B cells in mice boosted with all selected Ags except SA-Jo1 (**Supplementary Fig. 4D**). In addition, we have tested the frequency of the nucleosome-specific B cells in mice boosted with SA-nucleosomes conjugates. In accord with previous study, we found (based on the staining with fluorescent nucleosome tetramers) that about ∼1% of GC B cells in SA-DEL immunized mice boosted with SA were specific to nucleosomes (Gonzalez-Figueroa *et al*., 2021). However, boosting of mice with SA-nucleosomes led to significant drop in the frequency and numbers of the nucleosome-specific GC and memory cells (**Supplementary Fig. 4E-H**). Overall, this data suggests that nucleosomes and other NucPr Ags contain peptides cognate to Tfr cells and if acquired by GC B cells may promote their suppression by Tfrs.

### Formation of conjugates between human B cells acquiring NucPrs and Tfrs

While antigen-specific targeting of NucPrs to GC B cells promotes rapid accumulation of Tfr cells in mice, whether this finding maybe relevant to the regulation in humans is unclear. We therefore assessed whether human Tfrs may have a subset of cells cognate to the NucPrs. Based on the previous studies, cognate interactions between B and T cells cocultured *ex vivo* lead to increased formation of B-T cell conjugates detectable by flow cytometry analysis (Choudhuri et al., 2005). We therefore utilized a similar approach with Tfrs (see **Fig. 7A**) and B cells (CD19^+^ CD3^-^ CD27^-^) sorted from the blood of healthy human subjects. To promote antigen uptake by human B cells we first coupled anti-human IgM-biotin antibodies to SA, SA-DEL or SA-NucPrs to generate protein complexes αIgM (αIgM-SA or αIgM-SA-DEL) and αIgM-NucPrs (αIgM-SA-NucPr). Sorted B cells were then cultured in the presence of αIgM or αIgM-NucPrs to promote crosslinking of B cell receptors and internalization of the linked to αIgM protein complexes for degradation and HLA class II /peptide presentation. Tfrs from the same patient were then co-cultured with activated or control B cells for 36h and formation of Tfr-B cell conjugates was assessed by flow cytometry analysis (**Fig. 7B**). In 3 independent experiments we detected an increase in the frequency of Tfr-B cell conjugates when Tfr were cocultured with activated B cells that have internalized NucPrs as compared to control Ags or B cells that did not acquire Ags (**Fig. 7B, C**). Based on that, we suggest that a significant fraction of the circulating human Tfrs may be specific to the peptides from the NucPrs selected in this study.

**Figure 7.**
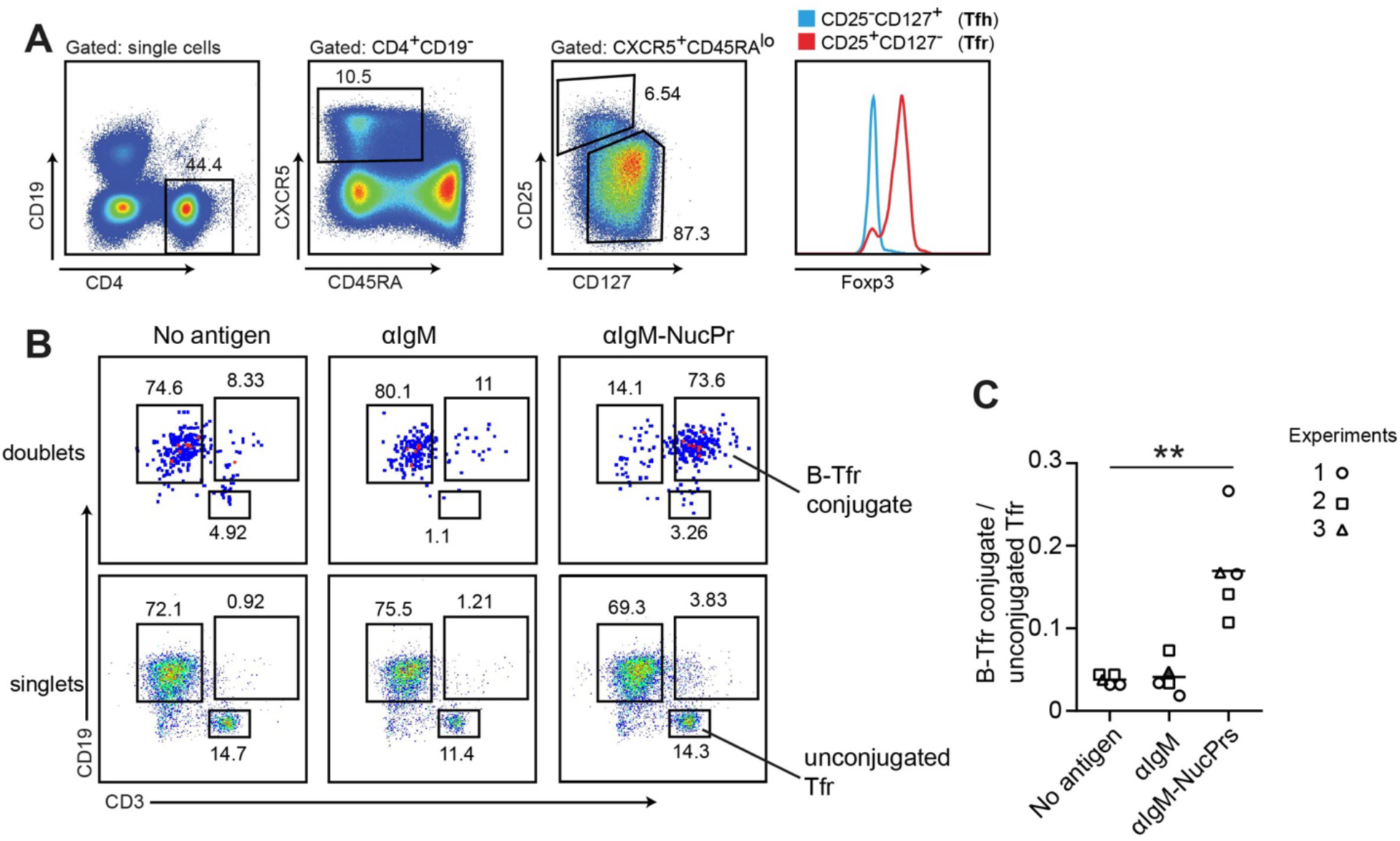
Formation of conjugates between human B cells acquiring NucPrs and Tfrs. B cells and Tfrs were sorted from human blood. (**A**) Sorting strategy for Tfr cells. (**B, C**) Sorted CD19^+^CD3^-^CD27^-^ naïve B cells were incubated ex vivo with αIgM (anti-IgM-SA (opened symbols), anti-IgM-SA-DEL (filled symbol)), αIgM-NucPrs (anti-IgM-SA-NucPrs) or medium for control for 30 min. After that they were cocultured with Tfrs from the same patient. (**B**) Representative example of flow cytometry analysis of singlets and doublets for B-Tfr conjugate formation after 36h of coculturing. (**C)** The ratio of Tfr-B cell conjugates (in doublets) to unbound Tfrs (in singlets). Data are from n=3 independent experiments. Each type of symbol represents one patient, 1 or 2 replicas. ** *p*<0.01. Two-tailed Student’s *t* test.

## Discussion

In this study we showed that combination of nuclear proteins (SS-A/Ro, RNP-Sm, Scl70, Jo-1 and nucleosomes) selected based on their frequent targeting by autoantibodies in autoimmune diseases, induces rapid accumulation of Tfr cells in the dLNs. While there is no detectable Tfr increase in mice directly immunized with NucPrs (conjugated to streptavidin and administered in Ribi adjuvant), a rapid Tfr response is detected when SA-NucPrs are administered to mice with an ongoing SA-specific GC response. In these mice, SA-specific GC (and possibly memory) B cells are expected to acquire SA-NucPrs via their B cell receptors, internalize the Ags and present both antigenic and NucPrs peptides in complex with MHCII for recognition by follicular T cells. While our study does not rule out the potential for dendritic cells and/or other Ag-presenting cells to promote Tfr responses at longer time scale, it suggests that direct targeting of selected NucPrs to abundant antigen-specific B cells via BCR could be utilized to trigger very rapid and robust increase in Tfr cells. Interestingly, a previous study suggested pausing of Tfrs in proximity of the tingible body macrophages that acquire dying B cells within GCs (Jacobsen et al., 2021). It is plausible to suggest that these macrophages may be another cell type in GCs that presents nuclear protein peptides for cognate stimulation of Tfr cells, possibly under all immunization conditions.

Based on the 10x genomics gene expression analysis, the majority of the NucPrs-induced Tfrs have distinct gene expression characteristics that suggest enhanced immunosuppressive functions. Based on the TCR immunorepertoire analysis, Tfr subset enriched in the SA-NucPrs boosted mice (cluster 10) have significant clonal overlap with other Tfr subsets. In contrast to this “immunosuppressive” subset, Tfr-like cell clusters 11 and 12 are predominantly overlapping with Tfh-like cell clones and may be related to the recently identified subset of Tfh cells in GCs that upregulate FoxP3 and play a role in the shutoff of GC response overtime (Jacobsen *et al*., 2021). Importantly, in mice boosted with SA-NucPrs we detected no increase in the Tfh-related clones in the “immunosuppressive”, as well as all other Tfr-like subsets. Therefore, we suggest that the majority of NucPrs-induced Tfrs are not derived from Tfh cells. Based on this, on the previously established overlap of Tfr and Treg TCR repertoire, as well as the important role of Ag-specific B cells in the very rapid induction (3 days) of the immunosuppressive Tfrs, we speculate that majority of NucPrs-induced Tfrs accumulate due to proliferation of preexisting Tfr clones in the follicles or their precursors at the T/B border following their cognate interactions with NucPr-acquiring and presenting B cells.

Identification of the dominant self-Ags cognate to Tregs remains an important task for both fundamental analysis of thymus-derived Tregs and for development of Tregs-dependent translational therapies (Leonard *et al*., 2017). Previous studies suggested that histones may contain Tregs epitopes. Vaccine therapy with peptides from nucleosomal histones H4 and H1 promoted Treg response, diminished auto-Ab levels and delayed the onset of nephritis in lupus-prone mice (Kang et al., 2011; Kang et al., 2005). However, availability of Tregs epitopes in the other NuclPr Ags has not been suggested before. Because we detected the strongest Tfr response after administration of the combination of NucPrs, we suggest that selected self-Ags contain multiple protein epitopes cognate to Tfrs. Importantly, our initial analysis of Tfrs isolated from human blood suggests that a fraction of these cells may be specific to the peptides from the NucPrs that we utilized in the study. Future studies should carefully delineate the peptide epitopes within nuclear proteins that induce Tfr responses in mice and humans and assess their potential for future vaccine therapies.

Our data suggest that Tfrs induced by SA-NucPrs boosting promote partial shut off of the overall GC response. Importantly, they more strongly impede SA-NucPrs-acquiring GC B cells and reduce their memory and antibody-secreting cell responses. Based on that, we conclude that cognate interactions with Tfrs promote superior suppression of GC B cells. The critical role of Tfrs in the observed regulation (rather than indirect effects from SA-NucPrs administration) is supported by complete reversal of the observed immunosuppression in the Tfr-deficient mice. To summarize, while out study does not rule out nonspecific inhibitory effect of Tfrs on non-cognate GC B cells, it suggests that GC B cells that acquire selected NucPrs will be subjected to specific repression by cognate Tfrs.

Multiple molecular mechanisms could potentially contribute to the negative control of GC B cells by cognate and noncognate Tfrs. NucPrs-induced Tfrs upregulate expression of multiple genes that may potentially contribute to the negative regulation of B cells and Tfh cells, including Ctla4, Icos, CD39, Granzyme B and Il10. Of note, expression of Neuritine that was recently shown to be produced by Tfrs and to limit B cell differentiation into plasma cells (Gonzalez-Figueroa *et al*., 2021), was not limited to the accumulating Tfr cluster with immunosuppressive phenotype. We speculate that multiple molecular mechanism may work in parallel to promote suppression of GC B cells. Cognate recognition by Tfrs of the GC-presented MHCII/peptides likely leads to more prolonged interactions between the cells (Jacobsen *et al*., 2021), increasing the duration of inhibitory signals received by GC B cells from cognate Tfrs. Future studies should dissect the contribution of distinct molecular pathways to the negative signals provided to GC B cells by Tfrs.

Future studies should also examine potential engagement of the NucPr-specific Tfrs for translational applications. To this end, our study suggests that SA-NucPrs boosting promotes a significant drop in the NucPrs-acquiring GC B cells not only in the wild-type, but also in the lupus-prone NZM2328 mice that have dysregulated GC responses and overtime develop anti-nuclear Abs. We therefore suggest that targeting NucPrs (and/or their specific peptides) to GC B cells should be further explored for analysis of their therapeutic potential for control of pathogenic B cells specific to autoantigens or/and allergens or promoting autoepitopes spreading for activation of pathogenic autoreactive T cells.

## Acknowledgements

We thank Prof. Jason Cyster (UCSF) for useful discussions and comments on the manuscript. Single cell processing and next-generation sequencing was carried out in the Advanced Genomics Core at the University of Michigan. Supported by the National Institute of Health R01 AI106806 and R21 AI142032 (IG), R01 AI132771 (ALD), R01-AR071384 (JMK), K24-AR076975 (JMK), P30-AR075043 (JMK, MPM).

## Author Contributions

Conceptualization, F.K, Z.B, and I.G.; Methodology, F.K., Z.B., A.L.D, J.M.K.; Investigation, F.K, Z.B, M.P.M, J.L.; Writing, F.K, Z.B. and I.G.; Funding Acquisition, I.G.; Supervision, I.G.

## Competing interests

JMK has received Grant support from Q32 Bio, Celgene/BMS, Ventus Therapeutics, and Janssen. JMK has served on advisory boards for AstraZeneca, Eli Lilly, GlaxoSmithKline, Bristol Myers Squibb, Avion Pharmaceuticals, Provention Bio, Aurinia Pharmaceuticals, Ventus Therapeutics, Vera Therapeutics, and Boehringer Ingelheim

## Methods

### Mice

C57BL/6 (B6, WT) mice were purchased from The Jackson Laboratory. Bcl6^fl/fl^ (Hollister *et al*., 2013) and Foxp3-YFP-cre mice (Rubtsov *et al*., 2008) were crossed in Alexander Dent’s Lab (Wu *et al*., 2016). NZM2328 mice (Jacob *et al*., 2003) were a gift to Michelle Kahlenberg from Chaim Jacob (USC). All mice were bred and maintained under specific pathogen-free conditions. Relevant mice were interbred to obtain Bcl6^fl/fl^ Foxp3-YFP-cre mice. All the animal experiments were conducted in compliance with the protocols reviewed and approved by the Institutional Animal Care and Use Committee of the University of Michigan.

### Antigen preparation

#### Duck egg lysozyme (DEL) purification

Duck eggs were locally purchased and DEL was purified as previously described (Allen et al., 2007). 400 mL of duck egg whites were blended with 2 L of buffer A (0.1M ammonium acetate buffer, pH = 9), filtered through 2 layers of Kim wipes, and stirred overnight (O/N) at 4°C with 4.5 g CM Sephadex C-25 beads (GE Healthcare) preequilibrated for 3 h with 100 mL of buffer A. The beads were then transferred into Buchner funnel covered with 7 layers of kimwipes, washed with 1 L of buffer A and eluted with 200-300 mL of buffer B (0.4 M ammonium carbonate buffer, pH = 9.2). The eluent was lyophilized, resuspended in 15 mL buffer A, and dialyzed two times in buffer A at 4°C. Insoluble material was removed by centrifugation for 5 minutes at 1000 rcf., and the supernatant was concentrated using a Centriprep filter (Millipore, MWCO 10,000 kDa) to 3 mL for loading onto a 1 × 66 cm gel filtration column comprised of 4 g of degassed Sephadex G-50 beads (GE Healthcare) preequilibrated in buffer A O/N. The column was washed with 200 mL of degassed buffer A and the concentrated eluent was then separated on the column using degassed buffer A to collect ten 5 mL fractions. Eluted fractions were analyzed by SDS-PAGE and stained with Coomassie brilliant blue. DEL-containing (band at ∼14 kDa) fractions were combined. DEL concentration was determined by SDS-PAGE using a standard curve with Hen egg lysozyme (HEL, Sigma).

### Generation of SA-DEL, SA-NucPr, and SA-OVA

SA-DEL was generated as previously described (Turner et al., 2017b). Purified DEL was conjugated to biotin at a 1:2 molar ratio using Sulfo-NHS-LC-Biotin (Thermo Fisher Scientific) according to the manufacturer’s directions and incubated for 3 hours on ice. After dialyzing 3 times in PBS DEL-biotin incubated with streptavidin (Sigma) at a 10:1 molar ratio for 30 minutes on ice, followed by removal of unbound DEL-bio by passage through a 30 kDa molecular weight cut-off desalting column (Bio-Rad). Nucleosome, RNP/Sm, Jo-1, Scl-70 and Ro (SSA) (AROTEC DIAGNOSTICS) were conjugated to biotin at a 1:50 molar ratio using Sulfo-NHS-LC-Biotin and incubated for 3 hours on ice. After dialyzing 3 times in PBS, Nucleosome-biotin, RNP/Sm-biotin, Jo-1-biotin, Scl-70-biotin and Ro (SSA)-biotin were incubated with Streptavidin at a 1:1 molar ratio for 30 mins on ice. Antigen conjugation was verified by SDS gel electrophoresis. After incubation, antigens were aliquoted and kept at -20°C. OVA-bio (Fisher, NC0887816) were incubated with Streptavidin at a 4:1 molar ratio for 30 mins on ice. All antigens were aliquoted and kept at -20°C.

### Generation of αIgM-SA, αIgM-SA-DEL, αIgM-SA-NucPrs

To generate anti-IgM-NucPr, Streptavidin, anti-IgM-bio, Nucleosome-bio, Jo-1-bio, Scl-70-bio, RO (SSA)-bio, and RNPsm-bio were incubated at a 1:2:0.5:0.5:0.5:0.5:0.5 molar ratio for 30 mins on ice. To generate anti-IgM-DEL, Streptavidin, anti-IgM-bio, and DEL-bio were incubated at a 1:2:2 molar ratio for 30 mins on ice. To generate anti-IgM-SA, Streptavidin and anti-IgM-bio were incubated at a 1:2 molar ratio for 30 mins on ice.

### Immunization

In some experiments mice were s.c. immunized with 50 µg SA-DEL, reimmunized with 10 µg SA, SA-OVA, or SA-NucPrs at day 8 after immunization. In some experiments serum from mice immunized with 50 µg SA at day 8 after immunization were collected and injected into unimmunized mice followed by immunization with 10 µg SA or SA-NucPrs. In some experiments naïve mice were immunized with 10 µg SA or SA-NucPrs directly. Lymphoid cells from the draining lymph nodes and the spleens of immunized mice were analyzed at the indicated time points. Blood was collected into Microvette CB 300 tube via cardiac puncture when the mice are under deep anesthesia. About 0.5 ml to 1 ml blood was obtained from one mouse. Serum recovered after centrifugation at 10000 g 5 min 20 °C.

### Flow cytometry analyses and FACS sorting

Single-cell suspensions from draining lymph nodes and spleens were prepared and filtered through a 70-µm nylon cell strainer (BD). Red blood cells were lysed. Cells were washed in FACS buffer (2% FBS, 1mM EDTA, 0.1% NaN_3_ in PBS) and followed by surface staining for the indicated markers for 20 min at 4 °C. NP-specific B cells or SA-specific B cells were detected with BCR specific binding with NP-PE (Biosearch Technologies) or SA-PE (BioLegend). For intracellularstaining, the FoxP3 intracellular staining kit (Thermo Fisher Scientific) was used according to the manufacturer’s instructions. Sample were then incubated with anti-FoxP3, CTLA4, Granzyme B, or Ig (H+L). All samples were acquired on a BD FACSCanto flow cytometer. For cell sorting, enriched B cells and T cells were incubated with antibodies in Sorting buffer (0.5% FBS and 2 mM EDTA in PBS) and were performed on a BD FACSAria III cell sorter. All data were analyzed with FlowJo (version 10.6.0) software.

### ELISA

SA-specific IgG were detected in serum from blood by ELISA. Nunc 96-well ELISA plates were coated with 50 µl of 2 µg/ml SA (Sigma) in borate saline buffer (100 mM boric acid, 0.9% NaCl, pH=7.4) overnight at 4 °C. Wells were blocked with 0.1% Biorad Gelatin in PBS plus 0.05% Tween-20. Twofold diluted serum sample were loaded into the plate. ELISA plates were incubated for 1 h at room temperature. Plates were washed with PBS containing 0.05% Tween-20. Bound antibody detected with 1.5 µg/ml IgG-HRP (Invitrogen). After washing, the color was developed with TMB (Thermo Fisher). The chromogenic reaction was stopped with 2N Sulfuric Acid and the plates were read with a Synergy HT microplate reader (Bio-Tek Incorporated) at 405 and 630 nm. All plates contained serial dilutions of the serum that was used to generate the calibration curve for quantitative comparison of the samples.

Anti-RNP was detected in female 52 week old C57Bl/6 and 32-52 week old NZM2328 mice (harvested at time of nephritis or at 52 weeks) via ELISA (Alpha Diagnostics, San Antonio, TX) according to manufacturer’s instructions.

### Immunofluorescence Staining

Freshly isolated lymph nodes were fixed in 1% PFA for 1 hour at room temperature, washed with PBS three times, and stored at 4ºC in 30% sucrose solution overnight. Fixed samples were then transferred to OCT (Tissue-Tek) and snap-frozen. 30 µm thick sections were cut via cryostat (Leica). The sections were dried at room temperature for 3 hours. They were blocked using normal rabbit serum (Sigma-Aldrich) in PBS with 0.2% Triton-X100 and then stained with anti-mouse CD3 PE-CF594 (BD) and anti-mouse Bcl6 Alexa Fluor 488 (BD) overnight. After washing, slides were then mounted in fluoromount-G (Southern Biotech) and analyzed via confocal microscopy with Leica SP5 II (Leica Microsystems) using a 25x objective. Images were processed using Imaris.

### 10x genomics and TCR immunorepertoire analysis

B6 mice were subcutaneously (s.c.) immunized with SA-DEL in Ribi and at day 8 were s.c. reimmunized with SA, SA-NucPr. Follicular T cells (CD4^+^PD1^hi^CXCR5^hi^) were sorted from mice at day 11 after immunization into PBS + 2% FBS. Single cell suspensions were subjected to counting and viability checks on the LUNA Fx7 Automated Cell Counter (Logos Biosystems) and diluted to a concentration of 700 -1000 cells/ul. Single cell libraries were generated using the 10x Genomics Chromium Controller with Immune Profiling reagents following the manufacturer’s protocol (10x Genomics). Final library quality was assessed using the LabChip GX (PerkinElmer). Libraries were subjected to paired-end sequencing according to the manufacturer’s protocol (Illumina NovaSeq 6000). Bcl2fastq2 Conversion Software (Illumina) was used to generate de-multiplexed Fastq files and the CellRanger Pipeline (10x Genomics) was used to align reads and generate count matrices. Barcodes with mitochondrial reads > 5% were removed. Preprocessing, clustering, and dimensionality reduction were performed using Loupe Browser and Loupe V(D)J browser (10x genomics, Cell Ranger). Graph-based clustering of Tfrs were visualized in 2D using UMAP algorithm.

For human scRNA-seq analysis, publicly available datasets were downloaded for lymph nodes cells from SARS-CoV2 mRNA-vaccinated patients (GSE195673) (Kim et al., 2022). Seurat v4 (Hao et al., 2021) was used for downstream analysis based on Seurat vignettes. Cells which had less than 200 or more than 8000 transcripts and mitochondrial genes representing greater than 15% of total transcripts were removed. All remaining cells were then clustered and projected into UMAP plots. The optimal number of PCs used for UMAP dimensionality reduction was determined using Jackstraw permutations which resulted in the first 15 PCAs being used. Identification of follicular T cells was accomplished by adding a module score in Seurat. Clusters in which the module score was significant enriched for were identified as being follicular. Follicular cells were then reclustered and differentially gene expression analysis between each cluster and all other cells was performed.

### Analysis of the human B and Tfr cells conjugate formation

Human Blood was collected with informed consent from healthy donors in accordance with a University of Michigan IRB approved protocol (HUM0007150). Fresh blood was subjected to centrifugation over a Ficoll-Paque Plus (GE Healthcare) density gradient, washed twice in PBS and resuspend cells in PBS. PBMC were stained with fluorophore-conjugated antibodies. After washing, CD3^-^CD19^+^CD27^-^ B cells and CD4^+^CD19^-^CD45RA^lo^CXCR5^+^CD25^hi^CD127^lo^ Tfr cells were sorted and resuspended in RPMI 1640 medium (GIBCO) with 10% FBS (Hyclone), 1% HEPES (Hyclone), 100 U/ml penicillin and 100 µg/ml streptomycin (GIBCO). 5×10^4^ B cells were first incubated with 5 µg/ml anti-IgM complexes and 1 µg/ml recombinant Human CD40L (Biolegend) for 30 mins and then 1×10^4^ Tfr cells were added. After incubation at 37°C and 5% CO_2_ for 36 hours, the cells were stained and analyzed by FACS. The ratio of CD3^+^CD19^+^ cells and CD3^+^ cells were calculated.

### Statistics

Statistical tests were performed as indicated using Prism 8 (GraphPad). No blinding or randomization was performed for animal experiments, and no animals or samples were excluded from analysis. All the statistical details of experiments and statistical analysis can be found in figure legends. Differences between groups not annotated by an asterisk did not reach statistical significance. Outliers were not excluded. Of note, outlier experimental data points in several figures did not affect statistical significance of the data.

| REAGENT or RESOURCE | SOURCE | IDENTIFIER |
| --- | --- | --- |
| **Antibodies** |  |  |
| Anti-mouse B220 Biotin (RA3-6B2) | BD Bioscience | Cat#553086 |
| Anti-mouse B220 Pacific Blue (RA3-6B2) | BioLegend | Cat#103227 |
| Anti-mouse B220 PerCP-Cy5.5 (RA3-6B2) | BD Bioscience | Cat#552771 |
| Anti-mouse B220 V500 (RA3-6B2) | BD Bioscience | Cat#561226 |
| Anti-mouse Bcl6 Alexa Fluor 488 (K112-91) | BD Bioscience | Cat#561524 |
| Anti-mouse CD3 Alexa Fluor 700 (17A2) | BioLegend | Cat#100216 |
| Anti-mouse CD3 PE-CF594 (145-2C11) | BD Bioscience | Cat#562286 |
| Anti-mouse CD4 APC/Cy7 (RM4-5) | BioLegend | Cat#100526 |
| Anti-mouse CD4 FITC (RM4-5) | BD Bioscience | Cat#553047 |
| Anti-mouse CD4 V500 (RM4-5) | BD Bioscience | Cat#560782 |
| Anti-mouse CD8a APC/Cy7 (53-6.7) | BioLegend | Cat#100714 |
| Anti-mouse CD8a Pacific Blue (53-6.7) | BioLegend | Cat#100725 |
| Anti-mouse CD8a V500 (53-6.7) | BD Bioscience | Cat#560776 |
| Anti-mouse CD8a Biotin (53-6.7) | BD Bioscience | Cat#553029 |
| Anti-mouse CD25 BV421 (PC61) | BioLegend | Cat#102034 |
| Anti-mouse CD38 PerCP-Cy5.5 (90/CD38) | BD Bioscience | Cat#562770 |
| Anti-mouse CD95 PE-Cy7 (Jo2) | BD Bioscience | Cat#557653 |
| Anti-mouse CD185 (CXCR5) BV605 (L138D7) | BioLegend | Cat#109110 |
| Anti-mouse CD279 (PD-1) PE-Cy7 (RMP1-30) | BioLegend | Cat#109110 |
| Anti-mouse Foxp3 APC (FJK-16s) | eBioscience | Cat#17-5773-82 |
| Anti-mouse GL7 eFluor 450 (GL-7) | eBioscience | Cat#48-5902-82 |
| Anti-mouse GL7 eFluor 660 (GL-7) | eBioscience | Cat#50-5902-82 |
| Anti-mouse Gr-1 Biotin (RB6-8C5) | BioLegend | Cat#108404 |
| Anti-mouse IgD APC/Cy7 (11-26c.2a) | BioLegend | Cat#405716 |
| Anti-mouse IgG HRP | Thermo Fisher Scientific | Cat#62-6520 |
| Anti-mouse Ig(H+L) Alexa Fluor 488 (Polyclonal) | Southern Biotech | Cat#1010-30 |
| Anti-human CD3 AF700 (OKT3) | BioLegend | Cat#317339 |
| Anti-human CD19 Pacific Blue (SJ25C1) | BioLegend | Cat#363036 |
| Anti-human FoxP3 AF488 (259D) | BioLegend | Cat#320211 |
| Anti-human CD4 BV510 (RPA-T4) | BioLegend | Cat#300545 |
| Anti-human CD127 BV605 (A019D5) | BioLegend | Cat#351333 |
| Anti-human CD25 PE (M-A251) | BioLegend | Cat#356103 |
| Anti-human CD45RA APC-Cy7 (HI100) | BioLegend | Cat#304127 |
| Anti-human CXCR5 AF647 (RF8B2) | BioLegend | Cat#558113 |
| Anti-human CD27 FITC (M-T271) | BD Bioscience | Cat#555440 |
| Anti-human IgM Biotin | Jackson Immuno Research Labs | Cat#109-066-129 |
| **Chemicals, peptides, and recombinant proteins** |  |  |
| Ficoll-Paque Plus | GE Healthcare | Cat#17-1440-02 |
| EZ-Link Sulfo-NHS-LC-Biotin | Thermo Fisher Scientific | Cat#A39257 |
| Fluoromount-G | SouthernBiotech | Cat#0100-01 |
| 1-step Ultra TMB-ELISA Substrate Solution | Thermo Fisher Scientific | Cat#34028 |
| Foxp3/Transcription Factor staining Buffer Set | eBioscience | Cat#00-5523-00 |
| Gelatin blocking buffer 1% in PBS | Alfa Aesar | Cat#J62755 |
| Recombinant Human CD40L | BioLegend | Cat#591702 |
| Ovalbumine | Sigma | Cat#A5503-5G |
| CM Sephadex C-25 beads | GE healthcare | Cat#17-0210-01 |
| Sephadex G-50 medium beads | GE healthcare | Cat#17-0043-01 |
| Sephadex G-100 beads | GE healthcare | Cat#17-0060-01 |
| Sigma Adjuvant System | Sigma | Cat#S6322-1VL |
| Streptavidin | Sigma | Cat#S4762-5MG |
| Streptavidin PE | BioLegend | Cat#405204 |
| Streptavidin Qdot 605 | Thermo Fisher Scientific | Cat#Q10101MP |
| Streptavidin Qdot 647 | Thermo Fisher Scientific | Cat#S21374 |
| Streptavidin Qdot 655 | Thermo Fisher Scientific | Cat#Q10121MP |
| Nucleosome | AROTEC DIAGNOSTICS | ATN02-02 |
| Jo-1 | AROTEC DIAGNOSTICS | ATJ01-02 |
| Scl-70 | AROTEC DIAGNOSTICS | ATS01-02 |
| Ro (SSA) | AROTEC DIAGNOSTICS | ATR02-02 |
| RNP-Sm | AROTEC DIAGNOSTICS | ATR01-02 |
| **Experimental Models: Organisms/Strains** |  |  |
| C57BL/6J | The Jackson Laboratory | JAX:000664 |
| Bcl6<sup>fl/fl</sup> | (Hollister et al., 2013) |  |
| Foxp3-YFP-cre | (Rubtsov et al., 2008) |  |
| NZM2328 | (Jacob et al., 2003) |  |
| **Software and Algorithms** |  |  |
| FlowJo V10 | FlowJo | https://www.flowjo.com/ |
| Graphpad Prism 8 | GraphPad | https://www.graphpad.com/ |
| Imaris | Bitplane | https://imaris.oxinst.com/ |
| Adobe Illustrator | Adobe | https://www.adobe.com/ |
| Loupe Browser 6.0 | 10x genomics, Cell Ranger | https://www.10xgenomics.com |
| Loupe VDJ Browser 4.0 | 10x genomics, Cell Ranger | https://www.10xgenomics.com |

---

**Supplementary Figure 1.**
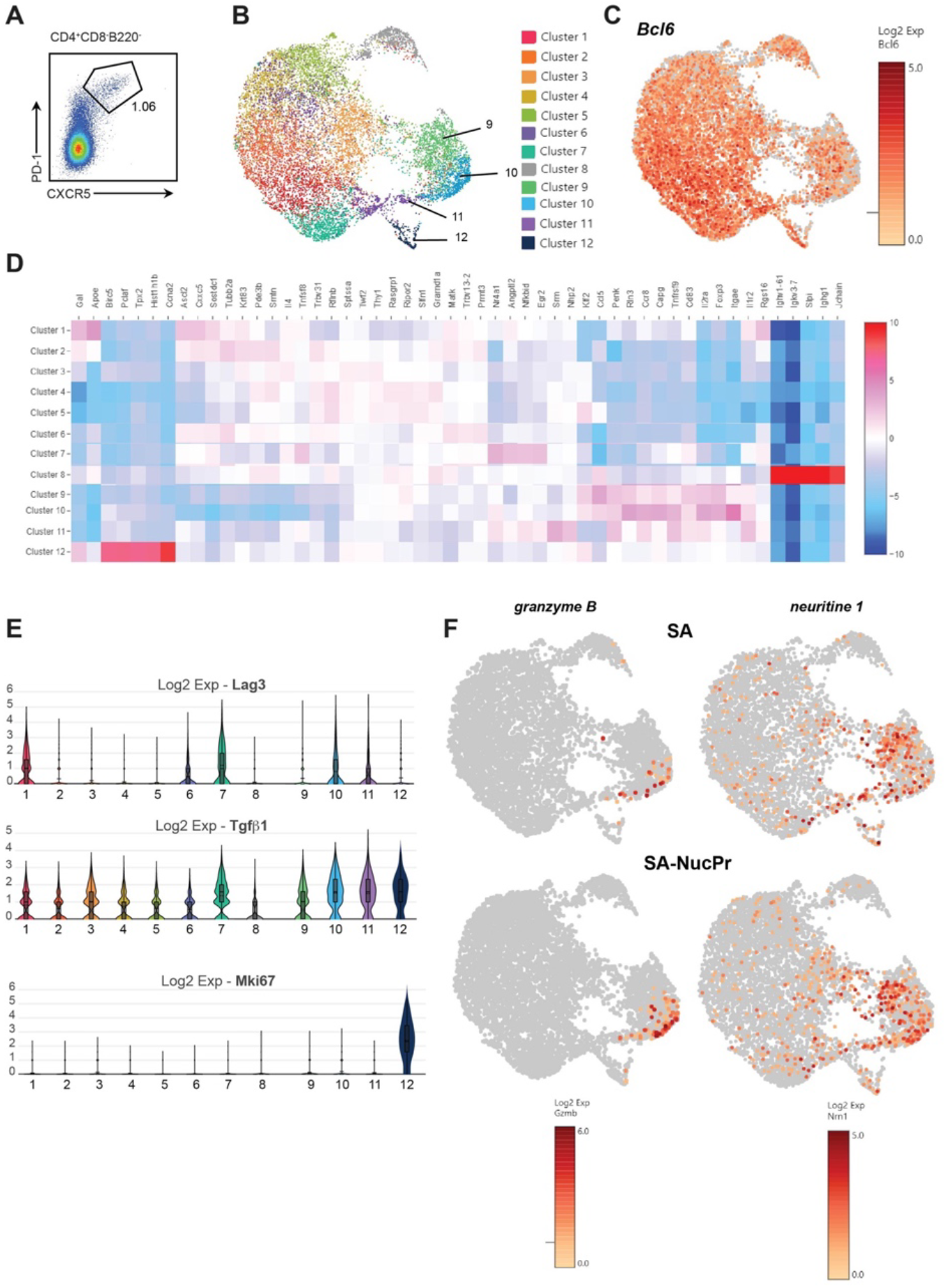
Analysis of Tfr cells gene expression and TCR repertoire in the SA and SA-NucPr-boosted mice. Single cell 10x genomics (Cell Ranger) analysis of gene expression of the CD4^+^CD8^-^B220^-^CXCR5^hi^PD1^hi^ cells (Tfr and Tfh) sorted from the dLNs of mice treated as described in **Fig. 1A** following their boosting with SA (2 mice) or SA-NucPr (2 mice). (**A**) The gating strategy for follicular T cell sorting. Cell purity > 94%. (**B**) Graph-based clustering of follicular T cells from combined SA and SA-NucPr boosted mice visualized in 2D using uniform manifold approximation and projection for dimension reduction (UMAP) algorithm. (**C**) Single cell expression of follicular T cells transcription factor *Bcl6*. (**D**) Heatmap with genes that have highest statistical significance of mean gene expression in Tfr clusters 1-12 (5 genes per each cluster) calculated based on negative binomial test (Cell Ranger). (**E**) Expression of selected genes associated with Treg-mediated regulation and proliferation in the follicular T cell clusters 1-12. (**F**) Single cell expression of granzyme B and neuritine1 in Tf cells.

**Supplementary Figure 2.**
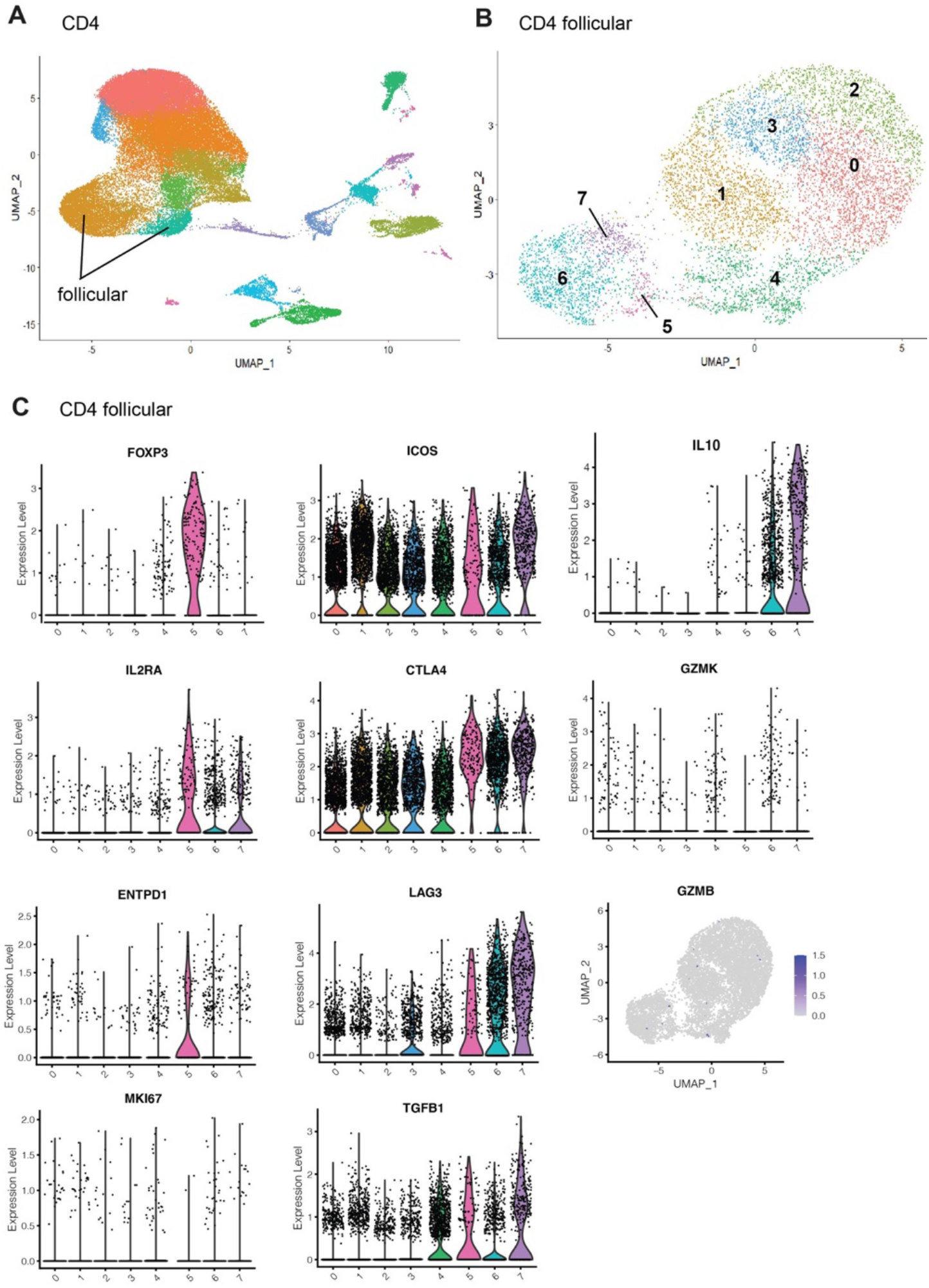
Analysis of human Tfr cells gene expression. Human lymph nodes scRNA-seq analysis was performed on the publicly available single cell datasets from fine needle aspirations of the draining ipsilateral axillary lymph nodes in SARS-CoV-2 mRNA-vaccinated human subjects (GSE195673). Seurat v4 was used to normalize, find variable features (2000), and to perform PCA analysis of CD4 T cells. Follicular T cells were identified using modular score generated based on the expression of CXCR5, PDCD1, BCL6, ICOS, CTLA4, IL1R2 and CXCL13. Follicular T cell subsets were then reclustered. (**A, B**) PCA and clusterization analysis for CD4 T cells (in **A**) and follicular-like CD4 T cells (in **B**). (**C**) Expression of the immunosuppressive genes and genes associated with cell proliferation in human follicular-like T cell clusters 0-7. Clusters 4-7 are more consistent with Tfr-like cells.

**Supplementary Figure 3.**
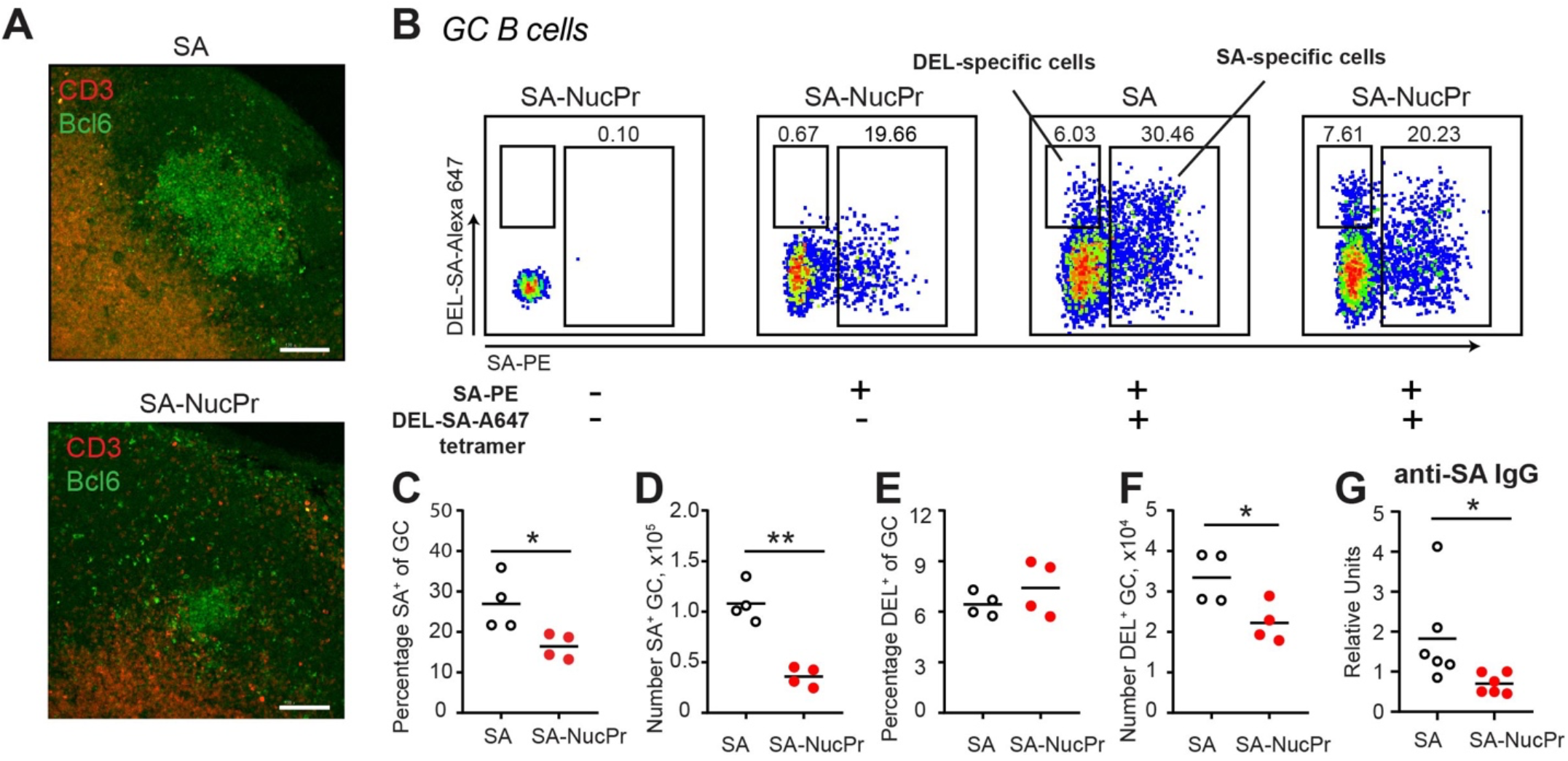
Boosting mice with SA-NucPrs suppresses GC, memory and PB responses with predominant inhibition of the SA-specific B cells (related to Fig. 3). (**A**) Immunofluorescent analysis of GC responses in the dLN sections of mice immunized and boosted with SA or SA-NucPr as in **Fig. 1A**. Scale Bar, 100 µm. Representative images from 2 independent experiments, dLNs from 8 mice. (**B-G**) Analysis of SA-specific and DEL-specific B cell responses in mice immunized with SA-DEL in Ribi s.c. and boosted with SA or SA-NucPr as in **Fig. 1A**. To identify DEL-specific and SA-specific GC B cells lymphocytes from dLNs were stained with SA-PE and DEL-SA-A647 tetramers. (**B**) Representative flow plots for DEL-specific and SA-specific (B220^+^CD3^-^FAS^+^GL7^+^IgD^lo^) GC B cells in SA- and SA-NucPr-boosted mice. (**C, D**) The frequency (**C**) and the number (**D**) of SA-specific GC B cells. (**E, F**) The frequency (**E**) and the number (**F**) of DEL-specific GC B cells. (**G**) Serum titers of SA-specific IgG in SA and SA-NucPr boosted mice. Data are from n=2 (**C-F**) or n=3 (**G**) independent experiments. Each symbol represents one mouse. Lines indicate means. * *p*<0.05, ** *p*<0.01, two-tailed Student’s T test.

**Supplementary Figure 4.**
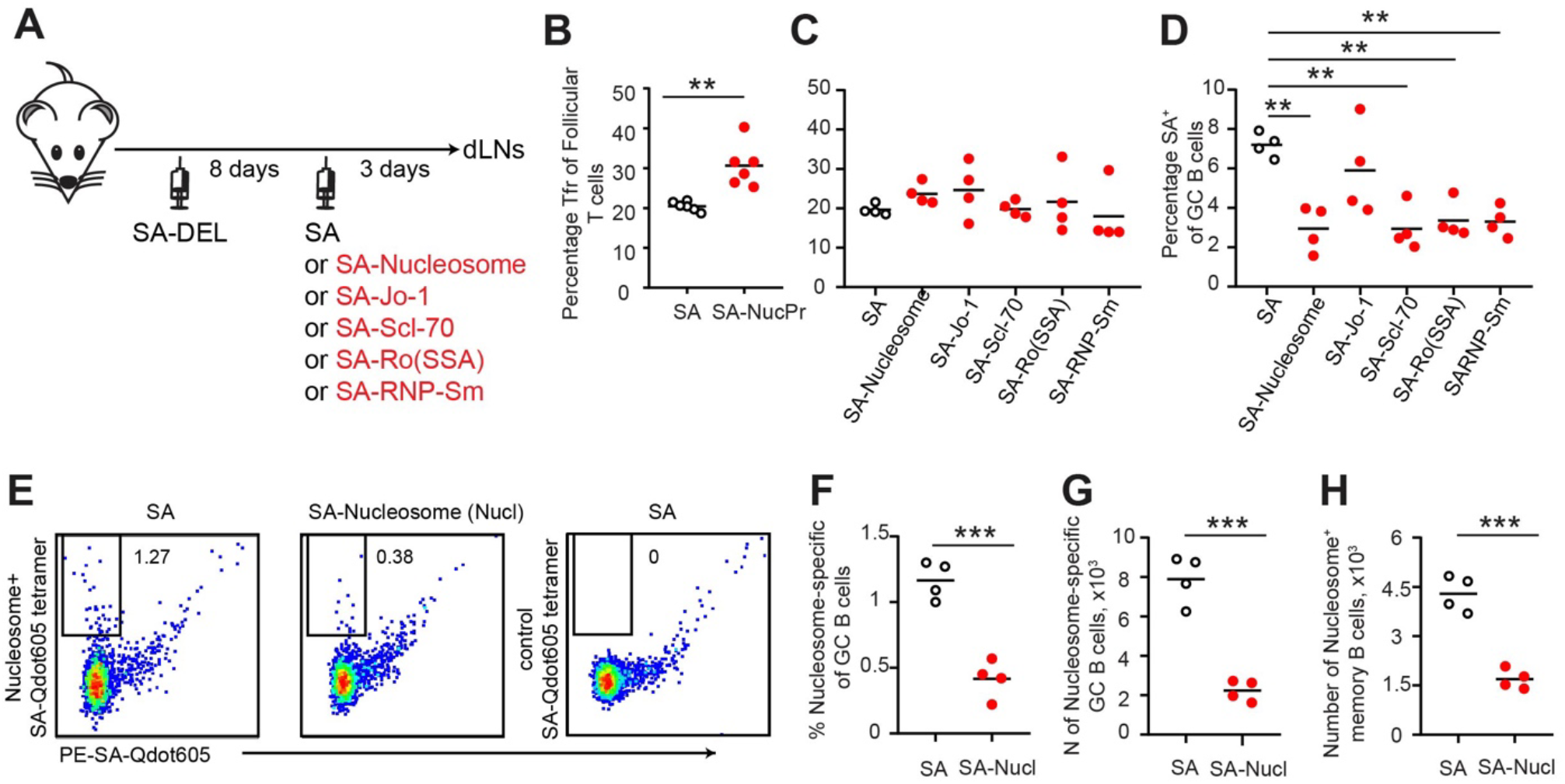
Analysis of Tfr and GC B cell responses in mice boosted with SA linked to individual NucPrs/complexes. (**A-D**) Analysis of Tfr and SA-specific GC response in B6 mice immunized with SA-DEL in Ribi s.c. and at day 8 s.c. reimmunized with SA, SA-Nucleosome, SA-Jo-1, SA-Scl-70, SA-Ro (SSA), or SA-RNP-Sm in Ribi for analysis 3 days later. (**A**) Experiment outline. (**B, C**) Tfr cells percentage of total CD4^+^B220^-^CXCR5^high^PD1^high^ follicular T cells in mice boosted with SA or combined SA-NucPrs (**B**) and in mice boosted with SA versus separate SA-NucPrs (**C**). (**D**) The SA-specific B cells percentage of total GC B cells. (**E-H**) Analysis of the nucleosome specific B cells in SA-DEL-immunized mice, boosted with SA or SA-nucleosomes. (**E**) Representative flow analysis of nucleosome-specific GC B cells. To identify Nucleosome-specific and SA-specific GC B cells lymphocytes from dLNs were stained with PE-SA-Qdot605 and Nucleosome-SA-A647 or control SA-A647 tetramers. (**F-H**) Nucleosome-specific GC B cells percentage of total GC B cells (**F**), GC B cell numbers (**G**), memory B cells (B220^+^CD3^-^FAS^lo^GL7^lo^CD38^hi^IgD^lo^) (**H**). Data are representative of n=2 independent experiments. Each symbol represents one mouse. Lines indicate means. ** *p*<0.01, *** *p*<0.001. Two-tailed Student’s *t* test (for **B, F, G, H**). One-way ANOVA analysis with Bonferroni’s multiple comparisons test (for **C, D**).

**Supplementary Table 1.** Significant genes in the clusters 1-12 of murine follicular T cells. Single cell 10x genomics (Cell Ranger) analysis of gene expression in FoxP3 expressing CD4^+^CD8^-^B220^-^CXCR5^hi^PD1^hi^ cells sorted from the dLNs of mice treated as described in **Fig. 1A**. Tfr cells were reclustered in Loupe software (Cell Ranger) based on gene expression into clusters 1-12 (see **Fig. 2**). Statistical significance of mean gene expression in Tfr cluster compared to other Tfr cells was calculated based on negative binomial test (Cell Ranger).

